# A recurrent sequencing artifact on Illumina sequencers with two-color fluorescent dye chemistry and its impact on somatic variant detection

**DOI:** 10.1101/2025.09.27.678978

**Authors:** Beverly J. Fu, Vinayak V Viswanadham, Dominika Maziec, Hu Jin, Peter J. Park

## Abstract

**Background:** The sequencing-by-synthesis technology by Illumina, Inc. enables efficient and scalable readouts of mutations from genomic data. To enhance sequencing speed and efficiency, Illumina has shifted from the four-color base calling chemistry of the HiSeq series to a two-color fluorescent dye chemistry in the NovaSeq series. Benchmarking sequencing artifacts due to biases in the newer chemistry is important to evaluate the quality of identified mutations.

**Results:** We re-analyzed a series of whole-genome sequencing experiments in which the same samples were sequenced on the NovaSeq 6000 (two-color) and HiSeq X10 (four-color) platforms by independent groups. In several samples, we observed a higher frequency of T-to-G and A-to-C substitutions (“T>G”) at the read level for NovaSeq 6000 versus HiSeq X10. As the per-base error rate is still low, the artifactual substitutions have a negligible effect in identifying germline or high variant allele frequency (VAF) somatic mutations. However, such errors can confound the detection of low-VAF somatic variants in high-depth sequencing samples, particularly in studies of mosaic mutations in normal tissues, where variants have low read support and are called without a matched normal. The artifactual T>G variant calls disproportionately occur at NT[TG] trinucleotides, and we leveraged this observation to bioinformatically reduce the T>G excess in somatic mutation callsets.

**Conclusions:** We identified a recurrent artifact specific to the Illumina two-color chemistry platform on the NovaSeq 6000 with the potential to contaminate low-VAF somatic mutation calls. Thus, an unexpected enrichment of T>G mutations in mosaicism studies warrants caution.

## BACKGROUND

The reversible terminator DNA sequencing chemistry^1^ underlying the Illumina sequencing instruments produces the vast majority of the world’s sequencing data. This chemistry permits “sequencing-by-synthesis” in which a DNA sequence is decoded through the iterative construction of a complementary strand. Each new base on this complementary strand is built from a deoxynucleotide triphosphate (dNTP) containing a fluorescent dye, which is imaged when the dNTP is successfully incorporated and before the dye is cleaved off to allow a new labeled dNTP to lengthen the strand.

Various aspects of this technology were refined over the years, enabling greater accuracy and throughput in terms of both read length and the number of fragments sequenced at once. Major platform updates included HiSeq 2000 in 2010, HiSeq X Ten (X10) in 2014, NovaSeq 6000 in 2017, and NovaSeq X in 2022.

The key difference between Illumina’s NovaSeq and HiSeq series lies in the number of distinct dye colors used to distinguish the four major types of dNTPs. HiSeq chemistry uses four fluorescent dyes to differentiate each nucleotide on a flow cell **(Figure 1)**. However, starting with NovaSeq 6000, Illumina adopted a two-dye system to differentiate each base: T by green, C by red, A by both red and green, and G by neither color. This binary color-coding system reduces both reagent consumption and the time required to sequence each base, as the instrument takes only two rather than four images of fluorescent dyes during each cycle. Despite these efficiencies, whether this simplified chemistry achieves the same or better base-calling accuracy across diverse sequencing contexts has not been thoroughly investigated.

**Figure 1:**
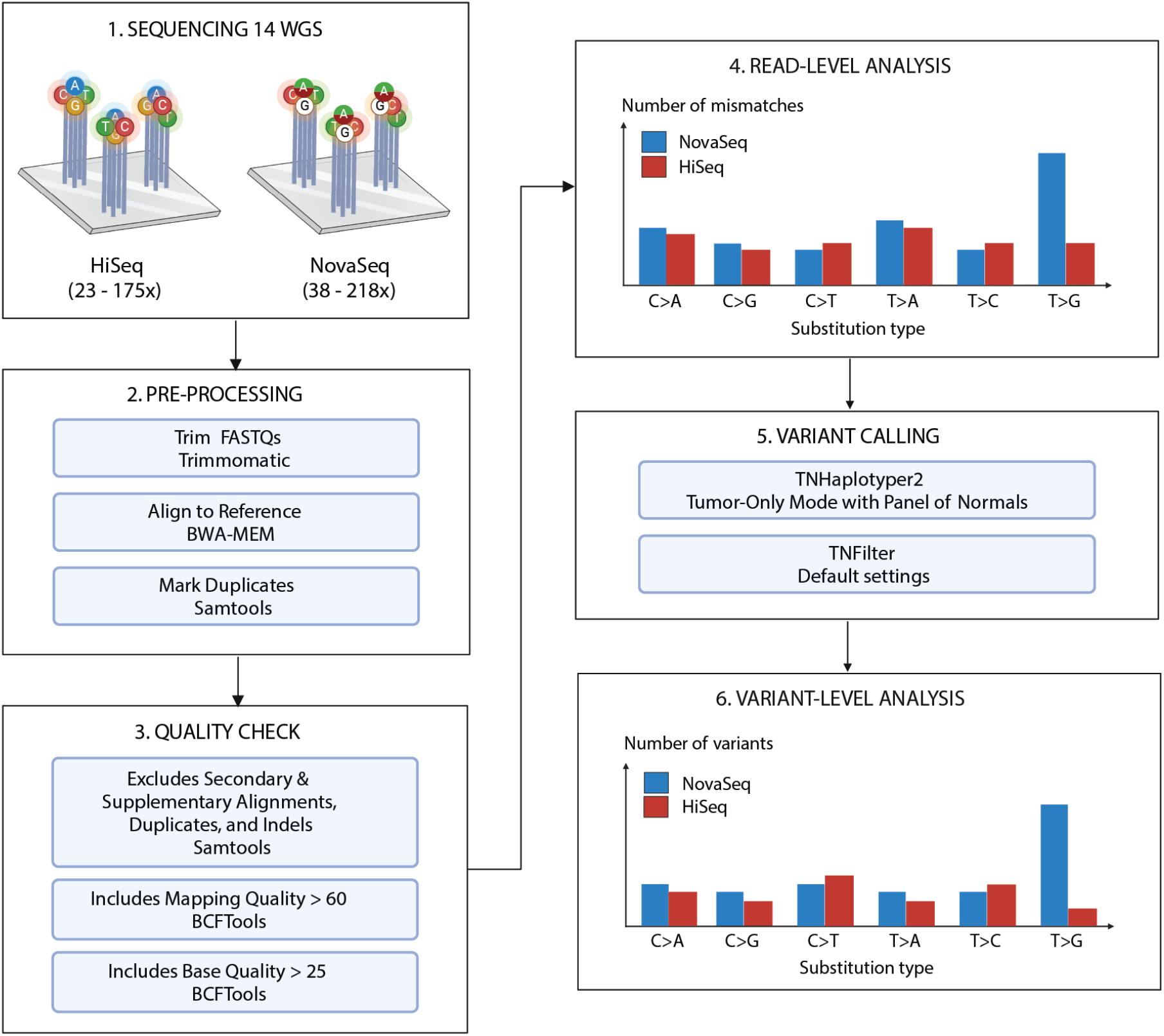
Overview of pre-processing and analysis steps. 14 WGS published datasets were compared, with the coverage ranges shown. The raw data were processed uniformly using standard processing and quality control steps. Both read-level and variant-level analyses comparing the four-color (HiSeq) and two-color (NovaSeq) dye platforms were performed. For variant calling, we used TNHaplotyper2, which is a faster reimplementation of MuTect2.

Systematic errors across generations of Illumina sequencers have been documented in the literature. For instance, patterns of decreasing base quality toward the end of reads have been observed from the very beginning^2^. For the two-color system, overcalling of G base (which is called when A/C/T signal is absent) has been observed^3^, because it is not possible to distinguish between the presence of G and the absence of any signal due to, e.g., degrading of the cluster or physical blocking of flow cell imaging. In an extreme manifestation of this error, a persistent loss of signal over multiple sequencing cycles leads to a “poly-G” sequence that spans up to the end of the read. Trimming poly-G’s from read ends is a typical bioinformatic solution to this problem. Otherwise, there is no clear consensus on how to distinguish true N>G mutations from subtler losses of signal using existing QC tools. One clever way to identify sequencing errors is to examine the overlapping region between the paired-end reads when the sequenced fragment is shorter than the length of both reads combined. Stoler and colleagues^4^ used this approach to comprehensively measure error rates and sequence contexts of errors in >1900 *E. coli* datasets from 7 Illumina platforms, finding lower error rates for more recent platforms. Stoler et al also determined that it is challenging to separate operator effect from systematic errors, as different groups have observed different sequencing outcomes despite using the same technology.

We became interested in systematically characterizing Illumina sequencing errors when we observed different frequencies of a specific substitution type in two Illumina instruments. In our study of mosaic mutations in human tissues (somatic variants in non-cancer tissues are often referred to as ‘mosaic’ variants), we had performed high-coverage (>200X) whole-genome sequencing (WGS).

Unexpectedly, we found a marked elevated frequency of T>G and A>C substitutions in samples sequenced using the NovaSeq 6000 platform. In our previous studies of mosaic mutations in the neurotypical brains sequenced on a HiSeq X10^5–7^, we had not observed such T>G enrichment (hereafter, we refer to T>G and A>C collectively as T>G unless indicated otherwise, since one cannot distinguish the strand in which the error occurred).

In the present manuscript, we sought to determine whether the enrichment of T>G could have been caused by platform-specific artifacts. We examined 14 pairs of publicly available human whole genome samples sequenced on both NovaSeq and HiSeq between 2012 and 2021 **(Materials and Methods; Supp. Table 1**). Our analysis shows evidence of excess T*>*G substitutions at the read level in 11 out of 14 whole genome samples sequenced on NovaSeq compared to HiSeq and that the severity of this excess varies across affected samples. Arora and colleagues^8^ already noted the existence of a higher frequency of T*>*G substitutions. Sequencing three cancer cell lines up to 278X on both platforms, Arora et al found that the NovaSeq 6000 produced more homopolymers of Gs and T>G mutations than HiSeq X. However, their report was limited to 3 cell lines sequenced at a single facility. Our study, on the other hand, conducts a more comprehensive analysis across 14 pairs of cell lines and primary tissue samples sequenced at several centers over many years. More importantly, our analysis reveals that the frequency of the artifactual mutation is sufficiently low so that standard analysis of germline (~50% allele fraction) variants or somatic variants with matched controls should not be affected under conventional sequencing depths. However, in situation where somatic variants of interest may have extremely low variant allele frequency (VAF)—such as variants that arise in resistance to therapy or in non-neoplastic tissues without matched controls—variant calling methods are applied with looser criteria to increase detection sensitivity. We show that it is important to be aware of the potential T>G artifacts in these studies.

## RESULTS

To investigate NovaSeq-specific artifacts, we re-analyzed 14 pairs of human WGS datasets, each consisting of one NovaSeq- and one HiSeq-generated dataset derived from the same biological sample or library. The sample set comprised four cancer cell lines (KM12, HCT116, U2OS, HCC1395), nine widely used non-cancer cell lines (HG001-HG007, NA12891, NA12892), and a brain sample (UMB1465). These datasets span a broad range of sequencing depths (23X-218X), were generated over 9 years on several versions of HiSeq (2000, 4000, X) and NovaSeq 6000, and were published in multiple independent studies ^7,9–12^. Notably, two sample pairs (UMB1465 and HCC1395) were sequenced to >100X on both platforms, allowing us to examine the presence of the artifacts that are unlikely to be observed in standard coverage data. The detailed metadata (including library preparation method, sample-specific coverage, and accession number) are provided in **Supp Table 1**.

### T>G mismatches are significantly elevated on NovaSeq at the read-level

First, we quantified the T>G mismatches at the read level using the 14 matched NovaSeq/HiSeq datasets (**Materials and Methods**). Each sample was pre-processed to minimize the presence of known sequencing artifacts common to both platforms, such as read-start contamination by adapters and read-end base quality degradation (**Figure 1; Materials and Methods**). Stringent base quality and alignment quality thresholds were applied to remove non-platform-specific artifacts and isolate NovaSeq-specific artifacts. In an example read-level view (**Figure 2A**), we can confirm that there are very few conventional mismatches due to sequencing or alignment errors. To quantify mismatch profiles across datasets, we examined platform-specific mismatches across the six categories of single nucleotide substitutions, collapsing reverse complements into unified categories (e.g., T>G and A>C are indistinguishable and are thus combined). A platform-specific mismatch was defined as a site where a mismatch is detected on NovaSeq but not HiSeq **(Figure 2A)**, or vice versa.

**Figure 2:**
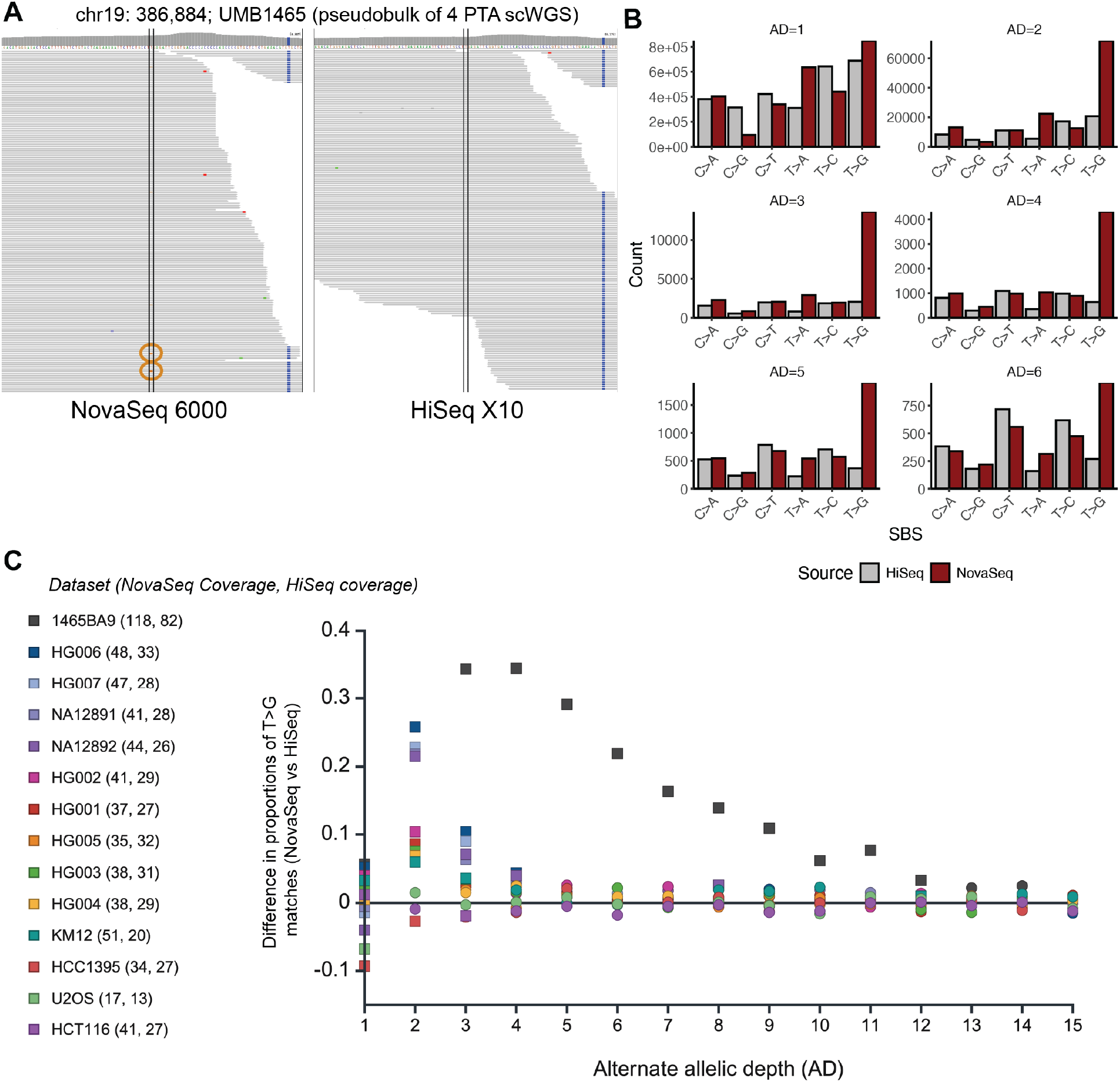
Pileups of T>G/A>C mismatches are significantly enriched in NovaSeq 6000 versus HiSeq X10 sequencing reads. (A) A browser view of a characteristic platform-specific enrichment event at chr19:386784 (delineated by the vertical dotted lines in each panel) within UMB1465-BA9. The reference (ref) allele is T, and the alternate (alt) allele G (circled in brown) is detected along 2 reads (AD=2) on NovaSeq (left), but 0 reads on HiSeq (right). (B) Number of mismatches (sites) at different allelic depths (ADs) for NovaSeq and HiSeq for UMB1465-BA9. Allelic depth (AD) ranges from 1 to 6. The results shown are for chr 19. (C) The differences in the proportion of T>G mismatches between NovaSeq and HiSeq. The numbers next to the sample names are the sequencing depths on NovaSeq and HiSeq. A difference greater than 0 indicates a T>G excess on NovaSeq. Squares instead of circles denote a significant difference in the proportion of T>G variants (α = 0.05) at that AD, based on a two-sided proportion Z-test.

We initially focused on sample UMB1465-BA9, derived from the cortical grey matter of a human donor. The WGS data from each platform for this sample is a pseudo-bulk formed by merging single-cell WGS datasets of 4 neurons, resulting in coverages of 218X for NovaSeq 6000 and 160X for HiSeq 2000. We measured the distribution of mismatches as a function of the allelic depth (AD) of the alternative allele (**Figure 2B**). At AD=1, the number of mismatches is high across all substitution types, with only mild enrichment for T>G. This suggests that, although we have removed a large fraction of sequencing/alignment errors with our stringent filtering, some errors remain. However, at ADs 2−5, the NovaSeq-sequenced dataset shows a pronounced excess of T>G compared to its HiSeq counterpart, with no comparable excess for any other single-base substitution (SBS) types (**Figure 2B**). Notably, at AD = 3, the NovaSeq dataset shows a 580% increase in T>G mismatched sites, with some secondary increases such as at T*>*A sites. At AD > 5, the NovaSeq excess starts to moderate, although the T>G frequency is still the highest at AD=6.

We expanded this analysis to the 13 other pairs to examine whether T>G excesses also contaminated other NovaSeq datasets. To account for inter-sample variation in sequencing depths, we measured the excess of T*>*G mismatches as a proportion of total mismatches detected at a specific allelic depth. For example, at AD = 3 in sample UMB1465-BA9 **(Figure 2C)**, T*>*Gs as a fraction of all single nucleotide mismatches collapsed into their reverse complements (C>A, C>G, C>T, T>A, T>C, T>G) are 0.57 on NovaSeq and 0.23 on HiSeq (**Supp Info 1**). We disregard significant read-level differences between the two platforms where AD = 1, as mismatches with such minimal read-support are unlikely to be called as variants. For each sample and AD, we tested the significance of the platform-wise difference in T*>*G proportions using a two-sided proportion Z-test (*α* = 0.05) under the null hypothesis that there would be no difference between the platforms (**Methods**). Based on this enrichment metric, 11 of 14 pairs showed significant enrichment for T*>*G mismatches in NovaSeq data at ADs between 2 and 10 **(Figure 2C)**. We observed stronger enrichments for samples at low ADs if the NovaSeq dataset was sequenced to deeper depths, although the enrichment was far more pronounced in the >100X UMB1465-BA9 pseudobulk. Collectively, these results demonstrate that read pileups in NovaSeq 6000 exhibit a disproportionate burden of T>G mismatches compared to HiSeq X10, supporting the presence of a platform-specific artifact.

### T>G mismatches are also significantly elevated on NovaSeq at the variant-level

We next investigated whether the enrichment of T>G mismatches observed at the read level in NovaSeq 6000 data propagates to somatic variant callsets. The presence of such mismatches in variant calls depends on whether the T>G substitutions pass all post-calling filters applied during somatic mutation analysis **(Supp. Table 2)**. To evaluate this, we generated somatic mutation callsets for each sample using TNHaplotyper2 (**Methods**). We compared the variant calls generated from matched NovaSeq 6000 and HiSeq X10 datasets of 10 of the 14 samples. We did not proceed with the pseudobulk from UMB1465 that we previously used for pileup analysis for the variant calling strategy. An appropriate somatic variant callset for any sample in our cohort should comprise both potential sequencing artifacts and legitimate somatic mutations at a broad range of allele frequencies. The true mutations from a pseudobulk of 4 30-60X single-cell WGS datasets would comprise only those at AFs of 25%, 50%, 75%, or 100% representing mutations found in 1, 2, 3 or all 4 cells; respectively. Any variant calls called from the pseudobulk with AFs significantly less than 25% would comprise only sequencing errors and no legitimate mutations, leading to a separate ascertainment bias in variant calling. Due to differences in sequencing depth on each platform yielding differing numbers of mutation calls, we computed the difference in the proportions of variant calls from each platform that were T>G. We evaluated the significance of this difference using a two-proportion z-test (**Methods**) at a significance level of p<0.05.

Examining variants identified exclusively on one platform, we found that 6 out of the 10 samples continued to display a significant excess of T*>*G on NovaSeq at AD ≤ 10, while 2 additional samples display a significant excess of T>Gs at ADs 10-20. The remaining 2 samples, HG002 and HG005, showed no T>G enrichments on NovaSeq over HiSeq. The 8 out of the 10 samples showing any enrichment showed sequencing depths of 30-47X; enrichment of T>G calls at AD=2 suggests that T>G mutation excesses in NovaSeq are found at minimum AFs of 4.25 %. In the sample with the strongest enrichment overall (HG003), T>Gs at AD=3 (AF of ~7.8%) comprised 10% more of the mutation calls on NovaSeq versus HiSeq. The difference in the proportion of T>Gs had random fluctuations of 1-2% at the high AD range, but at lower AD, several samples had more than 5% difference **(Figure 3A)**. The strongest differences occurred between AD 2-5, suggesting that the T>G excess is most pronounced for low-VAF variant calls.

**Figure 3:**
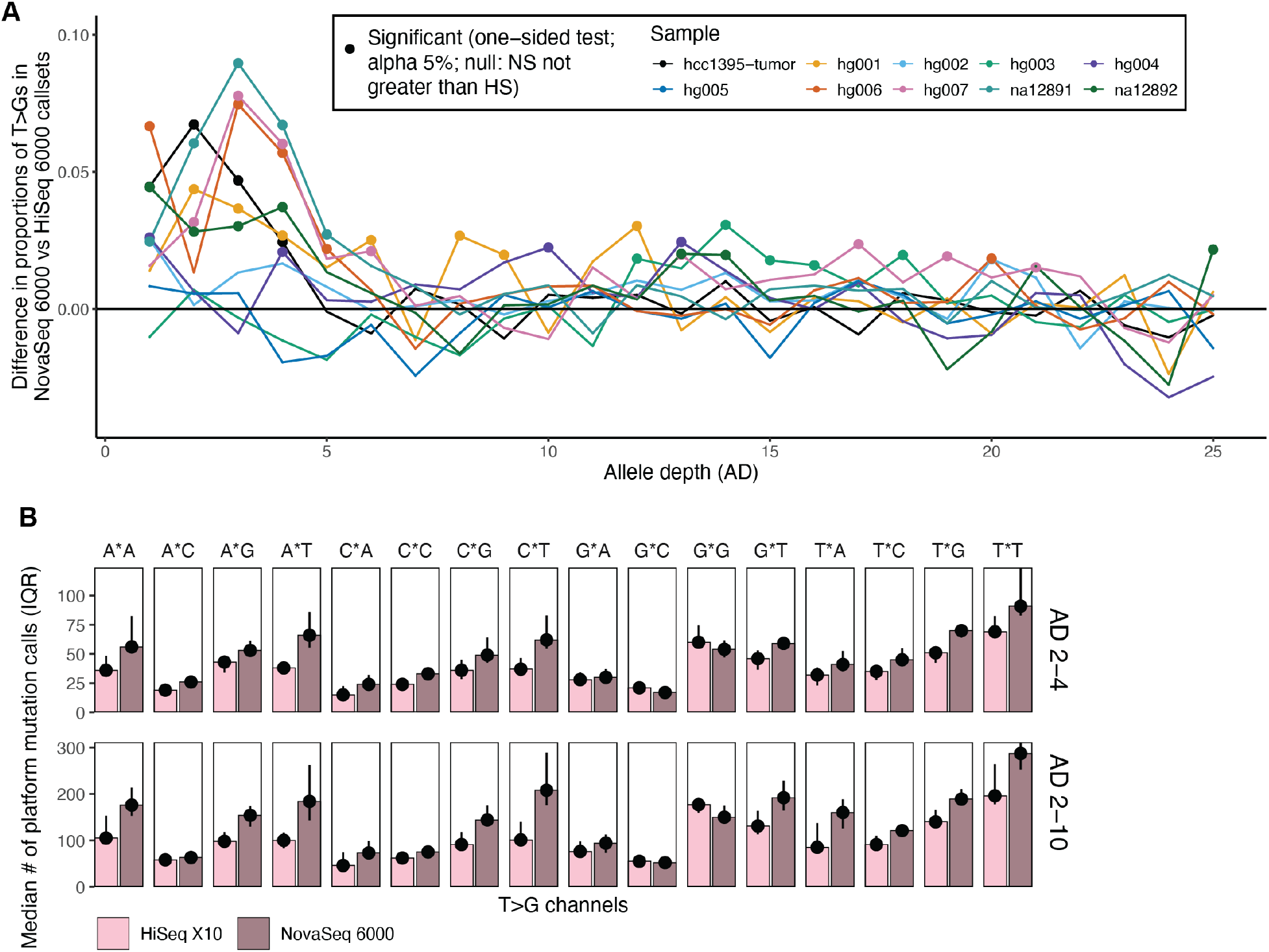
Enrichment of T>G mutations in somatic variant callsets on NovaSeq 6000. (A) Differences in the proportion of variant calls that are T>G/A>C at each allele depth (AD) in NovaSeq 6000 versus HiSeq X10. More positive values indicate greater enrichment for T>G mutations at that allele depth in NovaSeq 6000. Filled circles indicate a significant two-proportion Z-test result (*α =* 0.05) at that allele depth. (B) Trinucleotide spectra for NovaSeq 6000 and HiSeq X10 variant calls. Points and bars indicate the median number of mutations per platform per sample, while bars indicate the interquartile range (IQR). For each column, the “*” indicates where the T>G (COSMIC format) would occur (e.g. A*A would represent all ATA>AGA and TAT>TCT mutations).

### T>G mismatches on NovaSeq are enriched at NTT and NTG contexts

We hypothesized that if the NovaSeq-specific T>G excess appeared preferentially within specific genomic sequences, then it might be possible to filter out T>G mutations at those sequences without compromising entire variant callsets. Arora et al ^8^ previously reported that, even after filtering out low mapping quality (MQ<10) and low quality (BQ<10) bases, T>G mismatches remained enriched in the trinucleotide contexts of A[T>G]T, G[T>G]T, and T[T>G]T. To validate and extend these findings, we examined the nucleotide context of the excessive T*>*Gs on NovaSeq at the read level across multiple datasets. Specifically, we quantified the frequency of T*>*Gs at each of the 16 possible trinucleotide contexts N[T*>*G]N, following the COSMIC 96-channel format ^13^.

In NovaSeq datasets, T*>*Gs with ADs 2-4 were most frequently observed at N[T>G]T (A[T>G]T, C[T>G]T, G[T>G]T, and T[T>G]T) trinucleotide contexts (**Figure 3B**), although we observed several other contexts in which T>Gs were elevated in NovaSeq versus HiSeq. In particular, N[T>G]G contexts also showed notable excesses in NovaSeq over HiSeq. Although our variant-level enrichment analysis suggests that NovaSeq-specific enrichments were strongest between ADs 2-4, we also observed increases in NTT and NTG contexts overall between ADs 2-10 (**Figure 3B**). In contrast, HiSeq only showed consistent visible increases in T>Gs relative to NovaSeq for G[T>G]G contexts. Additionally, the overall distribution of trinucleotide contexts where T>Gs were discovered differed in the calls from each platform, with NovaSeq calls more elevated at NTT and NTG contexts overall.

Overall, we find that T>G mutations are moderately enriched in datasets generated from a NovaSeq 6000. This aligns with the observations of Arora et al ^8^, who found that T>G mismatches at the variant level persisted at [AGT]TT sequences and that using multiple callers could yield variants without the T>G excess. However, multi-caller strategies often show low sensitivities for low VAF variants, as not all callers are designed to detect those accurately^14^. Our harmonized analysis across many different independent samples establishes a more general bias towards T>Gs in N[T>G][TG] contexts across 10 biologically distinct samples sequenced on NovaSeq 6000. That such T>Gs occur consistently at low alternate allelic depths adds confidence to our hypothesis that the T*>*G mismatches detected on the samples are related to a persistent sequencing artifact.

### Sequence motif analysis removes excess T>G mismatches from standard variant calling workflows

The consistent trinucleotide signal suggests that mutational signature analysis could help remove the T>G excess in downstream analysis. First, to confirm the presence of the T>G artifact, we called somatic mutations using Mutect2 and MosaicForecast from newly collected deep WGS data (~210X) of 3 cerebral cortex samples each from 4 unrelated neurotypical individuals (i.e. 12 samples in all; **Supp Table 2**)^15^. Each sample was sequenced on either a HiSeq X10 or a NovaSeq 6000; none of the samples had paired datasets. Nevertheless, we found that the NovaSeq 6000-sequenced samples showed consistent and significant elevations of T>G mutations over HiSeq X10; we also observed smaller but still significant increases in T>C mutations (**Figure 4A**). T>Gs from NovaSeq 6000 were found at visibly-higher VAFs of 2-5% compared to 1-3% for HiSeq X10, while other mutations were found at comparable VAFs between the two platforms (**Figure 4B**). Mutations at VAFs of 2-5% for NovaSeq 6000 SBS’s suggest ADs ~4-10, consistent with the range observed for the T>G excess in our paired NovaSeq 6000/HiSeq X10 samples (**Figure 3A**). Thus, the T>G mutation bias is found with consistent properties in new and deeply sequenced samples.

**Figure 4:**
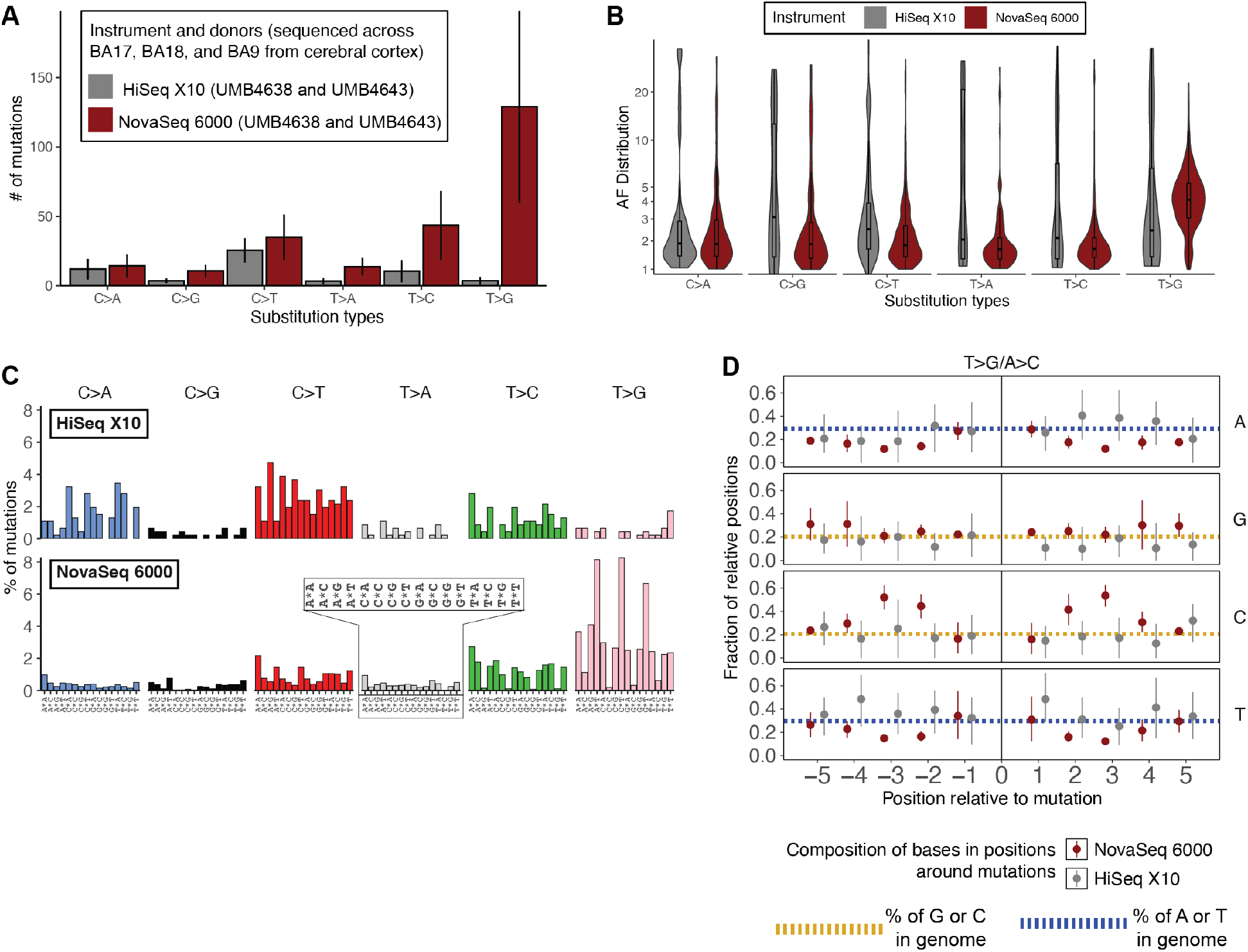
Sequence profile of NovaSeq-associated T>G mosaic variant calls in deep bulk whole-genome sequencing of non-cancerous human tissue. (A) Number of single-base substitutions (SBS’s) per COSMIC SBS type per platform. The line range shows 1 standard deviation of the average from 6 210X datasets per platform. (B) Allele fraction distributions for mutations per SBS per platform. (C) Trinucleotide spectra of SBS’s from samples sequenced on HiSeq X10 versus NovaSeq 6000. The trinucleotide channels (as in **Figure 3B**) are indicated in the inset. (D) Base composition in 10-base windows centered on NovaSeq 6000 SBS calls. At each relative position around all mutations (“0”) observed on each platform, we evaluated the fraction of bases that were each nucleotide (A, G, C, or T). Points represent the average fraction of bases containing a particular nucleotide, and line ranges show the 95% confidence interval. Gold and blue dotted lines show the genome-wide average of each nucleotide (gold for C/G and blue for A/T).

We found that mutations from NovaSeq 6000 exhibited very different sequence contexts compared to those from HiSeq X10. The trinucleotide spectra suggest that N[T>G]T mutations disproportionately make up all of the mutations called from NovaSeq 6000 (**Figure 4C**). The HiSeq X10 mutations resemble a signature more typically seen within clonal somatic mutations from noncancerous tissue ^5^. Examining broader sequences, we observed that NovaSeq 6000 T>G mutations disproportionately contained C/G bases within 5 bases of the mutation position (**Figure 4D**). On the other hand, T>A/A>T and T>C/A>G mutations in NovaSeq 6000 had similar surrounding base compositions as HiSeq X10 mutations **(Supp. Figure 1)**. The top motif near NovaSeq 6000 T>G mutations is a [T>G]TCC context, whereas G/A tetranucleotides dominate around A>C mutations (**Supp. Table 4**). Thus, we concluded that NovaSeq 6000 T>G mutations exhibit a distinct sequence profile compared to their HiSeq X10 counterparts.

Given the observed sequence differences around T>G mutations, we reasoned that the local sequence context from this set of samples should help filter out artifacts. We designed a strategy (**Figure 5A**) in which we computed the position weighted matrix (PWM) from the 8-base windows around somatic mutations from each platform in the unmatched samples. For HiSeq-generated WGS, MosaicForecast yields somatic mutations with a high validation rate of ~80% ^16^, and we obtained similar validation rates for many non-T>G mutations from NovaSeq 6000 ^15^. Thus, we hypothesized that the

**Figure 5:**
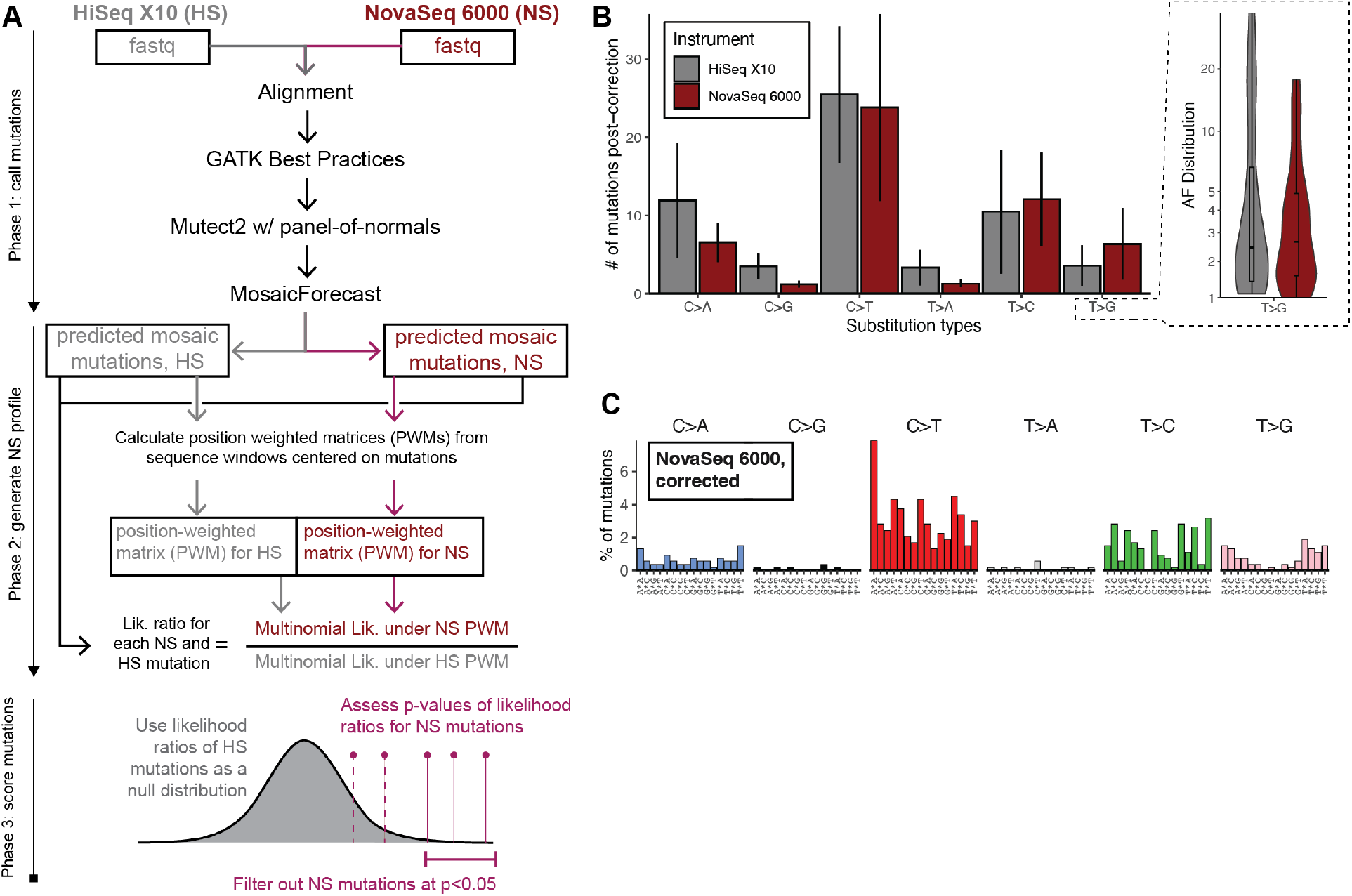
Correction of inflated somatic mutation counts using NovaSeq-specific sequence biases. (A) The workflow designed to correct NovaSeq 6000 callsets based on base composition as inferred from **Figure 4D**. (B) *Left panel:* Number of SBS’s per COSMIC SBS type per platform after correction. The line range is as in **Figure 4A**. *Right panel*: AF distributions of HiSeq X10 versus corrected NovaSeq 6000. (C) Trinucleotide spectrum of corrected NovaSeq 6000 counts.

PWMs of non-T>G mutations should reflect the true genomic background. Thus, true mutations should be similarly likely under the PWM from HiSeq X10 as NovaSeq 6000. Any deviations in the PWMs for T>G mutations between NovaSeq 6000 and HiSeq X10 should reflect artifact-prone genomic locations, and T>G mutations from NovaSeq 6000 would be significantly less likely to arise from the HiSeq X10 PWM. We calculated the likelihood ratio of any mutation arising from the NovaSeq 6000 PWM versus the HiSeq X10 PWM, and mutations significantly more likely to arise from NovaSeq 6000 (p<0.05) were marked as artifacts and filtered out.

The correction strategy significantly reduced the number of T>G’s from NovaSeq 6000 samples to comparable levels as HiSeq X10 while retaining similar numbers of non-T>G/A>C mutations (**Figure 5B, left panel**). In addition to achieving a significant, 25-fold reduction of T>G calls from NovaSeq 6000, the approach also rescaled the number of T>C calls (approximately 5-fold higher in NovaSeq 6000) to levels within 1 standard deviation of the HiSeq X10-generated calls. The AF distribution of the corrected NovaSeq 6000 mutations more closely matches that of the original HiSeq X10 calls (**Figure 5B, right panel**). Moreover, the trinucleotide spectrum of the corrected NovaSeq 6000 mutations more closely resembles that of the HiSeq X10 calls (**Figure 5C**). Overall, our workflow helps flag and correct T>G callsets down to appropriate sizes while retaining the properties of the likely true somatic mutations.

## DISCUSSION

Building on prior work,^8^ our findings establish that the NovaSeq-specific T>G excess is reproducible and recurrent across independently generated datasets, produced at multiple institutions over many years. The samples with the artifacts spanned diverse samples, from non-tumor samples (e.g., neurotypical brain sample 1465BA9) to gold-standard Genome in a Bottle samples^12^ (HG001-HG007).This consistency suggests that the artifact represents an inherent issue with sequencing on the NovaSeq 6000 instrument.

We hypothesize that the artifact tends to occur in 2-5% of sequenced bases, particularly at NT[TG] sites. In genomic libraries sequenced at lower depths (<30X), excess T>G mismatches would appear primarily at AD < 2, insufficient for most algorithms to call as a variant. However, at higher depths (>50X), the mismatches may occur at AD ≥ 2, overlapping the ADs of the rarest true mosaic mutations discoverable in such datasets. Thus, although its low rate of occurrence has made it nearly undetectable in conventional germline studies, the T>G artifact may cause confound the detection of true mutations in mosaicism studies and other settings in which efforts are made to identify very low VAF variants. For example, the artifact would pose challenges for analyzing normal tissues, where most somatic mutations occur at low VAFs and do not climb to the kinds of VAFs seen in driver or passenger mutations in cancer. With deeper sequencing efforts being undertaken by consortia such as the Somatic Mosaicism Across Human Tissues Network ^17^ to study somatic variation in noncancerous tissue, one must pay careful attention to systemic artifacts such as the one we describe here.

The distribution of trinucleotide contexts for artifactual N[T>G]T closely mimics those of known COSMIC somatic mutational signatures SBS9 (associated with polymerase eta), SBS17b, and SBS28.

Although the biological etiologies of SBS17b and SBS28 are uncertain, SBS17b is currently attributed to the chemotherapeutic 5-fluorouracil ^18^ and to damage by reactive oxidative species. Our samples—mostly non-tumor samples not exposed to 5-fluorouracil—exhibit T>G enrichment only when sequenced on a NovaSeq, effectively excluding SBS17b as the underlying cause (SBS28 resembles SBS17b and its etiology is unknown). However, its resemblance to a COSMIC signature makes it difficult to find by standard mutational signature analysis in cases whether SBS17b cannot be excluded *a priori*. Despite the evidence that we and others have presented about the existence of a NovaSeq artifact, we cannot validate its true mechanism without access to Illumina’s proprietary chemistry. We hypothesize that the binary two-dye fluorescent system makes it difficult to distinguish loss of-signal events during base detection from true G nucleotides. A loss of signal may occur for various reasons, including cluster degradation or physical blockage of the flow cell by particulates during the imaging process.

A major contribution of this manuscript is the proposed method for ameliorating the impact of the artifact. Although trinucleotide context by itself cannot necessarily distinguish the artifacts from true mutations, we have proposed a PWM-based likelihood ratio method that utilizes a wider genomic sequence context. This approach allowed us to recover the correct trinucleotide spectra (that obtained from the HiSeq data) as well as expected frequency of T>G’s. We constructed PWMs by comparing NovaSeq and HiSeq datasets derived from entirely different samples and donors sequenced at different times but with otherwise similar properties (normal tissue and similar sequencing depths). The success of our approach suggests that similar PWMs could be constructed from the vast amount of HiSeq and NovaSeq data deposited in public genomic repositories.

Although more costly and technically challenging, an even better approach to deal with the T>G artifacts would be to employ newer experimental techniques. For example, error-correcting requires that reads originating from the same original DNA molecule or strand carry the same base mismatch. Unique molecular identifiers (UMIs) or distinct tags for the Watson and Crick strands of the same molecule (as used in duplex sequencing) mark the provenance of individual sequencing reads. In the absence of a true mutation, machine error is unlikely to generate the exact same mismatch on many independently sequenced reads from the exact same molecule and mapped to the exact same position. Thus, a mismatch would be retained for downstream bioinformatic analysis only if a sufficiently large set of reads with the same UMIs or strand tags unanimously support the mismatch.

## METHODS

### Read-level processing

To compare mismatch profiles at low allele depths (ADs) of the two platforms with greater confidence, we applied stringent pre-processing steps to remove platform-agnostic artifacts or errors associated with low read quality. First, we applied Trimmomatic^19^ to remove the final 5 bp of each read to account for read-end base quality degradation. We also trimmed the first 8 bp (for 101bp reads) or 12 bp (for 151bp or 250-bp reads, as in the KM12 and HCT116 samples, respectively) of each read to remove possible read-start contamination due to adapter sequences. To further ensure that only high-quality reads were retained, we computed the Phred-scaled average base quality within 10-bp sliding windows across each read. We made a cut at the leftmost position of the window if the Phred-scaled average base quality of the assessed window dropped below 25 and discarded the rest of the downstream read sequence. We retained only those reads possessing a minimum number of bases after trimming (65 for 101 bp paired-end reads, and 100 for 151/250 bp paired-end reads respectively). This approach is more stringent than typical read preprocessing done in advance of variant calling and was only taken to control for sequencing noise when examining T>G pileups.

Following realignment of the remaining reads, we filtered duplicate alignments. We further removed reads if (1) they were not mapped in proper pairs (e.g., correct orientation and both pairs mapped), (2) they had supplementary alignments, (3) they possessed a Phred-scaled mapping quality score of less than 60, or (4) their CIGAR strings reported indel or clipping operations. We used the final set of aligned reads to assess T>G mismatches in our sequencing datasets.

### Mutation calling

To prepare variant calls from the 14 NovaSeq/HiSeq sample pairs’ WGS data (**Supp Table 1**), we applied the GATK Best Practices to preprocess BAMs without the extensive read-level processing applied for pileup analysis. We re-aligned and harmonized all processing steps to the GRCh38 reference genome (“GCA_000001405.15_GRCh38_no_alt_analysis_set.fna.gz” from NCBI). As we could not infer if BQSR was uniformly applied to each dataset, and as we wanted to preserve the base qualities as originally reported with published and available datasets, we did not apply BQSR ourselves in downstream processing. Sentieon’s TNHaplotyper2^20^ was applied to the 14 NovaSeq/HiSeq sample pairs in tumor-only mode with a panel of normals (Mutect2-WGS-panel-b38) for each sample and platform.

TNHaplotyper2 algorithm re-implements the mathematical models of the standard MuTect2, improving its computational efficiency without compromising the accuracy or consistency of its calls^20^. Thereafter, we applied TNFilter using its default settings (**Supp Table 2**) and examined only mutations that passed all filters. We then recalculated the raw alt allele depth (“alt AD”) for each variant call and retained those calls with alt AD ≥2. Finally, at each alt AD ranging from 1 to 25, we examined the differences in the numbers of T>G variant calls across platforms per sample (see below).

We also studied an additional six datasets generated from only a NovaSeq 6000: three cortical brain regions (BA9, BA18, and BA17) each taken from neurotypical individuals UMB5575 and UMB5580 (**Supp Table 3**). We identified mutations by first applying the standard GATK Best Practices to produce alignments to GRCh37d5 (including BQSR) followed by Mutect2. The mutation calls were then further refined using MosaicForecast, a method we developed that leverages read-level phasing and features to accurately distinguish mosaic SNVs from mutation call sets without the need for matched controls^16^. The MosaicForecast calls were compared to published mutation counts and profiles from two other neurotypical individuals, UMB4638 and UMB4643 ^5,15^, from whom deep WGS of the same cortical regions was sequenced on a HiSeqX10 only.

### Evaluating enrichment of T>G pileups and variant calls in NovaSeq 6000 versus HiSeq X10

To evaluate the enrichment of T>Gs in read-level pileups (**Figure 2C**) or variant calls (**Figure 3**), we implemented the two-proportion z-test. For each sample and each platform, we obtained the fraction of all pileups that are either T>G or A>C. We also computed the pooled fraction of T>G or A>C pileups by dividing the total number of A>C/T>G calls across both platforms by the total number of pileups. We then computed the z-statistic as

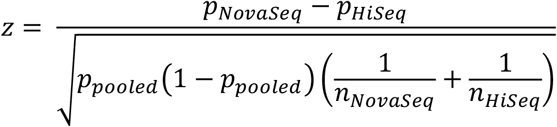

where *p*_*Novaseq*_ and *p*_*HiSeq*_ are the fractions of T>G/A>C pileups in NovaSeq and in HiSeq, respectively; *p*_*pooled*_ is the pooled fraction, and *n*_*Novaseq*_ and *n*_*HiSeq*_ are the total number of mismatched pileups in NovaSeq and in HiSeq, respectively. We used the same z-statistic for the variant calls and evaluated the statistical significance of the z-statistics for pileups and variant calls at α = 0.05.

## DECLARATIONS

### Ethics approval and consent to participate

Not applicable.

### Data Availability

The WGS data from the Broad Institute’s Cancer Cell Line Encyclopedia project (2012), the National Institute of Health (2020), and the Children’s Medical Research Institute (2021) were downloaded from the National Center for Biotechnology Information at https://www.ncbi.nlm.nih.gov/ using accession codes listed in **Supp. Table 1**. The WGS data from the National Institute of Standards and Technology’s Genome in a Bottle project (2012) was downloaded from the Google Brain Genomics Sequencing Dataset for Benchmarking and Development, publicly available at https://console.cloud.google.com/storage/browser/brain-genomics-public/research/sequencing;tab=objects?prefix=&forceOnObjectsSortingFiltering=false. Filters applied were “grch37” > “bam” > “novaseq” or “hiseqx” > “wgs_pcr_free” > “50x”. The WGS data from Harvard’s Brain Somatic Mosaicism Network (2017) with details in **Supp. Table 3** can be accessed from Synapse (https://www.synapse.org) under the ID syn21884970.

### Code Availability

The code for correcting the NovaSeq6000 mutation biases can be found at https://github.com/parklab/novaseq6000-correction

### Funding

This work was funded by Harvard Undergraduate Research Fellowship to BJF, R01CA269805 and R01HG012573 to PJP, and F31CA264958 to VVV.

### Competing Interests

The authors have no conflict of interest.

### Authors’ contributions

BJF, VVV, HJ, and PJP conceived the study. BJF and DM conducted upstream processing of the cell line and GIAB data with assistance from HJ. BJF conducted pileup analysis on the cell line and GIAB data with assistance from HJ. DM conducted variant calling on the cell line and GIAB data. VVV conducted variant calling on the cerebral cortex samples. VVV designed and implemented the mutation filtering strategy. BJF, VVV, and PJP wrote the manuscript with input from all authors. PJP supervised the study.

## Acknowledgements

The authors wish to acknowledge Michele Berselli for assistance with upstream bioinformatic processing of the GIAB samples, Lovelace J. Luquette for input on the read pileup analysis, and Sonia N. Kim and Sattar Khoshkhoo for helpful discussions.

## Notes

### Competing Interest Statement

The authors have declared no competing interest.

## REFERENCES

1. Bentley, D. R. et al. Accurate whole human genome sequencing using reversible terminator chemistry. Nature 456, 53–59 (2008).

2. Guo, Y., Ye, F., Sheng, Q., Clark, T. & Samuels, D. C. Three-stage quality control strategies for DNA re-sequencing data. Brief. Bioinform. 15, 879–889 (2014).

3. QC Fail Sequencing » Illumina 2 colour chemistry can overcall high confidence G bases. https://sequencing.qcfail.com/articles/illumina-2-colour-chemistry-can-overcall-high-confidence-g-bases/ (2016).

4. Stoler, N. & Nekrutenko, A. Sequencing error profiles of Illumina sequencing instruments. NAR Genom Bioinform 3, lqab019 (2021).

5. Bizzotto, S. et al. Landmarks of human embryonic development inscribed in somatic mutations. Science 371, 1249–1253 (2021).

6. Lodato, M. A. et al. Aging and neurodegeneration are associated with increased mutations in single human neurons. Science 359, 555–559 (2018).

7. Lodato, M. A. et al. Somatic mutation in single human neurons tracks developmental and transcriptional history. Science 350, 94–98 (2015).

8. Arora, K. et al. Deep whole-genome sequencing of 3 cancer cell lines on 2 sequencing platforms. Sci. Rep. 9, 19123 (2019).

9. Barroso-González, J. et al. Anti-recombination function of MutSα restricts telomere extension by ALT-associated homology-directed repair. Cell Rep 37, 110088 (2021).

10. Ghandi, M. et al. Next-generation characterization of the Cancer Cell Line Encyclopedia. Nature (2019) doi:10.1038/s41586-019-1186-3.

11. van Wietmarschen, N. et al. Repeat expansions confer WRN dependence in microsatellite-unstable cancers. Nature 586, 292–298 (2020).

12. Zook, J. M. et al. Extensive sequencing of seven human genomes to characterize benchmark reference materials. Sci Data 3, 160025 (2016).

13. Alexandrov, L. B. et al. The repertoire of mutational signatures in human cancer. Nature 578, 94–101 (2020).

14. Wang, Y. et al. Comprehensive identification of somatic nucleotide variants in human brain tissue. Genome Biology 22, 92 (2021).

15. Kim, S. N. et al. Cell lineage analysis with somatic mutations reveals late divergence of neuronal cell types and cortical areas in human cerebral cortex. bioRxiv 2023.11.06.565899 (2023) doi:10.1101/2023.11.06.565899.

16. Dou, Y. et al. Accurate detection of mosaic variants in sequencing data without matched controls. Nat. Biotechnol. (2020) doi:10.1038/s41587-019-0368-8.

17. Harris, E. NIH Launches Collaborative “Somatic Mosaicism” Network. JAMA 329, 1908 (2023).

18. Secrier, M. et al. Mutational signatures in esophageal adenocarcinoma define etiologically distinct subgroups with therapeutic relevance. Nat. Genet. 48, 1131–1141 (2016).

19. Bolger, A. M., Lohse, M. & Usadel, B. Trimmomatic: a flexible trimmer for Illumina sequence data. Bioinformatics 30, 2114–2120 (2014).

20. Kendig, K. I. et al. Sentieon DNASeq Variant Calling Workflow Demonstrates Strong Computational Performance and Accuracy. Front. Genet. 10, 736 (2019).

