## Supplemental Text for "A recurrent sequencing artifact on Illumina sequencers with two-color fluorescent dye chemistry and its impact on somatic variant detection"

**Supplementary Material**

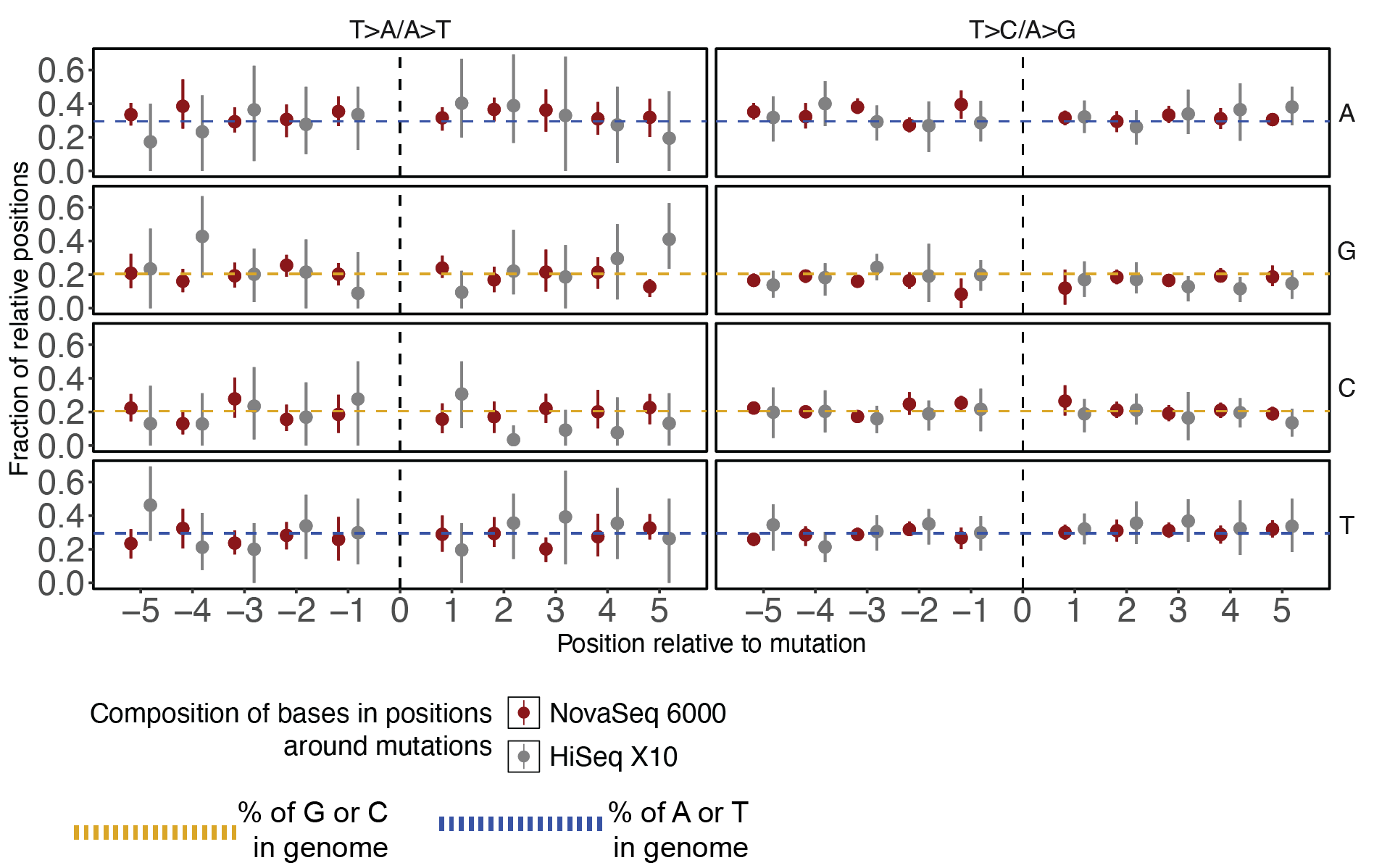

### Supplementary Figure 1

Base composition surrounding T>A/A>T and T>C/A>G mutations in the six platform-unmatched deep (>200X) human cortical samples (UMB4638, UMB4643, UMB5575, UMB5580; BA9, BA17, and BA18). The figure is laid out as in **Figure 4D.**

### Supplementary Information 1

Tables of the counts and proportions of all pileups of single-base substitutions (C>A, C>G, C>T, T>A, T>G, T>G) at different allele depths (AD) in the 14 different samples assessed for Figure 2. The difference in proportions between different platforms is given, along with a summary of proportions over different AD ranges for AD > 10.

### Supplementary Table 1

Metadata of 14 pairs of samples sequenced on both NovaSeq and HiSeq. The “Reference” column shows the reference genome to which the original dataset was aligned. For the variant-calling analysis, all samples were aligned to GRCh38. The “post-processing coverage” column indicates the coverage after applying the stringent read processing prior to substitution pileup analysis (**Figure 2**).

### Supplementary Table 2

List of filters applied during TNFilter to process mutation calls for the variant-level analysis (**Figure 3**).

### Supplementary Table 3

Metadata of deeply sequenced (>200X) human cortical samples. “Post-Procesing Coverage” refers to the BAM coverage after applying GATK4 Best Practices, not the read pileup pre-processing as in **Supplementary Table 1** or **Figure 2**.

### Supplementary Table 4

Summary of the k-mer (k=4) sequences found in the 20-base windows around different single-base substitutions discovered in the human cortical samples.
